# Genetics and genomics of social behaviour in a chicken model

**DOI:** 10.1101/165167

**Authors:** M. Johnsson, R. Henriksen, J. Fogelholm, A. Höglund, P. Jensen, D. Wright

## Abstract

The identification of genes affecting behaviour can be problematic, yet their identification allows a raft of possibilities. Sociality and social behaviour can have multiple definitions, though at its core it is the desire to seek contact with con- or hetero-specifics. The identification of genes affecting sociality can therefore give insights into the maintenance and establishment of sociality. In this study we used the combination of an advanced intercross between wild and domestic chickens with a combined QTL and eQTL genetical genomics approach to identify genes for social reinstatement (SR) behaviour. A total of 24 SR QTL were identified and overlaid with over 600 eQTL obtained from the same birds using hypothalamus tissue. Correlations between overlapping QTL and eQTL indicated 5 strong candidate genes, with the gene *TTRAP* being strongly significantly correlated with multiple aspects of SR behaviour, as well as possessing a highly significant eQTL. The distribution of eQTL can also indicate the genetic mechanisms underlying domestication itself. Multiple eQTL were found to in discrete clusters, however tests for pleiotropy show that these blocks were primarily linked in origin. This suggests that clustered genetic modules, rather than pure pleiotropy (as hypothesised by the neural crest theory) appears to be driving domestication in the chicken.

## INTRODUCTION

The identification of the genes responsible for behavioural variation has been an enduring goal in biology, with wide-scale ramifications, ranging from medical research to evolutionary theory on personality syndromes. For example, the identification of genes responsible for small-scale behavioural variation would more easily enable us to understand the genetic basis of personality behavioural disorders such as schizophrenia and bipolar disorder, as well shedding light on the types of genes involved, the selection acting on behavioural genes and how they can cause behavioural syndromes. While Quantitative Trait Loci (QTL) mapping has been successful in finding QTL for numerous behavioural traits, it has been considerably more difficult to isolate the actual genes underlying these QTL (reviewed by (FLINT 2003)). The same problems that limit other quantitative trait types are still a problem here (large confidence intervals generated in QTL analyses, epistasis within the QTL cross, QTL splitting down into multiple loci, etc), but are exacerbated by a phenotypic trait type that is often harder to define, ascertain and measure, making behavioural Quantitative Trait Gene (QTG) identification even more challenging.

Sociality and the genetics underlying it have been studied primarily in mammals (DONALDSON AND YOUNG 2008; MCGRAW AND YOUNG 2010; PERSSON *et al.* 2016), fish (BURMEISTER *et al.* 2005; WRIGHT *et al.* 2006a; WRIGHT *et al.* 2006b; GREENWOOD *et al.* 2016) and social insects (BEN-SHAHAR *et al.* 2002; ZAYED AND ROBINSON 2012). Sociality and social behaviour can have multiple definitions (BLEAKLEY *et al.* 2010), and can range from communication behaviour to the interactions between individuals of the same (and even different) species. For example dog (*Canis familiaris*) behaviour to seek out human contact and support (PERSSON *et al.* 2015), and honeybee foraging strategies and nursing behaviour are both forms of social behaviour. It is extremely diverse behavioural category, and can involve the production and reception of a variety of signals that go onto influence behaviour. A few specific genes have been found to determine certain aspects of sociality, for example *egr1* and song recognition in zebrafinch (MELLO *et al.* 1992), vasopressin (*v1aR*) expression in in voles, the *forager* gene (*for*) gene and its role in foraging strategies in different castes in the honeybee (BEN-SHAHAR *et al.* 2002).

Chickens are a social species that typically live in groups of between 6-10 individuals in the wild, that display a range of social behaviours(JOHNSON 1963). Social reinstatement, the desire of an animal to seek out conspecifics, is both a sociality and anxiety-related behaviour (MILLS AND FAURE 1991). It has been classically used in a variety of bird species to measure the strength of sociality at a base level, typically using a runway or treadmill test to assess an animal’s social motivation (SUAREZ AND GALLUP 1983b; JONES *et al.* 1991). Given the links to anxiety, the question of how social reinstatement (SR) relates to sociality in general is pertinent. In the case of the social reinstatement assay, perhaps the largest body of works concerns two lines of Japanese quail selected for high and low social reinstatement (SR). These birds have then been assessed for a wide variety of social assays to assess the extent to which selection for SR can affect other aspects of sociality. Launay et al(LAUNAY *et al.* 1991) found that high SR birds spent longer associating with conspecifics when given a paired goal box (one box empty, one box containing conspecifics). Two more studies found that when looking at pairs of high SR individuals in an open field arena they had a significantly shorter inter-individual distance as compared to the low SR birds (MILLS *et al.* 1992). High SR birds will even associate with conspecifics at the expense of food and water access, whilst high SR birds will also use more social facilitation by being able to learn to eat a novel food source by copying a conspecific ‘teacher’ (MILLS *et al.* 1997). More recently, high SR birds show a non-specific attraction for social conspecifics(SCHWEITZER *et al.* 2009), and have a consistently stable emotional reactivity, even in the face of high social instability(SCHWEITZER AND ARNOULD 2010).

Gene expression evidence in QTG identification in general is used when a candidate gene has been identified (see (WATANABE *et al.* 2007; LE BIHAN-DUVAL *et al.* 2011). Typically it is used to show differential expression in a tissue between cases and controls (DUBOIS *et al.* 2010). QTL for gene expression, termed genetical genomics or eQTL analysis, has also been used to try and identify quantitative trait genes (GILAD *et al.* 2008). In this instance, whole genome expression is combined with marker analysis to identify genes that are differentially expressed between the founder populations of the QTL cross, with these then being candidates for causative genes, especially if they overlap with existing QTL for the trait in question. An issue with this approach is that several eQTL may underlie a phenotypic QTL, and there is no way of further refining the selection to separate linkage from causation if they are performed on separate mapping populations. However, by fully integrating QTL and eQTL analysis in the same cross, it is possible to not only identify eQTL underlying a phenotypic QTL, but also to correlate actual gene expression with the desired trait in these overlapping loci. This can then rapidly identify quantitative trait genes that exhibit a close correlation with the phenotypic trait analysed and that underlie a particular QTL. This approach is obviously costly in terms of the number of individuals required for the eQTL analysis, but it has been used to identify QTG for lipid metabolism in mice (LEDUC *et al.* 2011) and startle-induced locomotion behaviour (amongst other ecological phenotypes) in *Drosophila* (AYROLES *et al.* 2009).

In this study, we identify a number of putative quantitative trait genes underlying phenotypic differences in social behaviour between Red Junglefowl and domesticated White Leghorn chickens. The domestic chicken exhibits a wide-range of behavioural as well as morphological differences, as compared to its wild-derived progenitor, the Red Junglefowl. This includes anxiety and sociality(SCHÜTZ *et al.* 2001), but also feather pecking behaviour and sociality(JENSEN AND WRIGHT 2014). Previous behaviour QTL work in the chicken has included mapping of feather-pecking behaviour (BUITENHUIS *et al.* 2003) and open field behaviour (BUITENHUIS *et al.* 2004) in layer chickens as well as fear-related behaviours and feeding behaviour in the F_2_ generation of the intercross line used in this work (SCHÜTZ AND JENSEN 2001; SCHÜTZ 2002; SCHUTZ *et al.* 2002; SCHUTZ *et al.* 2004). The study presented here continues from this using an advanced intercross (AIL) to generate small confidence intervals for mapping (DARVASI 1998). This AIL has already been used to identify genes affecting anxiety-related behaviour in an open field test (Johnsson et al 2016), however here we expand this to look at sociality. It was performed in three distinct phases. In the first phase, we performed QTL mapping of a social behaviour (n=572) as measured by a social reinstatement test in an 8^th^ generation advanced intercross of Red Junglefowl x White Leghorn chickens. In the second phase, a subset of the cross tested for behaviour (n=129) which has already been used in an expression QTL (eQTL) mapping of hypothalamic gene expression using a 135k probe array for each of these individuals, was used to overlap the previously detected eQTL with the newly detected behavioural QTL. Any gene that overlapped with a behavioural QTL was then checked for a correlation between gene expression and the behavioural trait in question and finally used in causation analysis using a Network Edge Orientation approach. In addition to this, the gene expression data was also correlated directly with the sociality phenotypes collected. This allowed us to assess genes with a high correlation with the behaviours, regardless of location, and also assess gene regulatory networks involving all those genes that were correlated with behaviour. Lack of extensive pleiotropic QTL and major trans-eQTL hotspots, and the presence of clusters of eQTL and behaviour QTL suggest a modular genetic architecture of chicken domestication, with the fine-scale mosaic of the advanced intercross also enabling the dissection of the hotspots that do occur to check if these represent truly pleiotropic loci.

## METHODS

### Chicken Study population and cross design

The intercross population used in this study was an eighth generation intercross between a line of selected White Leghorn (WL) chickens maintained from the 1960s and a population of Red Junglefowl (RJF) originally from Thailand (SCHUTZ *et al.* 2002; SCHUTZ *et al.* 2004). One male RJF and three female WL were used to found the intercross and generate 41 F_1_ progeny. The intercross was maintained at a population size of approximately 100 birds per generation until the F_7_ generation. The F_2_ intercross has previously been used to identify QTL for a number of different behavioural, morphological and life history traits (SCHUTZ *et al.* 2002; KERJE *et al.* 2003; WRIGHT *et al.* 2008; WRIGHT *et al.* 2010; WRIGHT *et al.* 2012). A total of 572 F_8_ individuals were generated from118 families using 122 F_7_ individuals (63 females and 59 males) and assayed for sociality behaviour. Average family size was 4.76 +/-3.1 (mean, s.d.) in the F_8_. 129 of the 572 F_8_ were used in an eQTL experiment, with the hypothalamus/thalamus dissected out at 212 days of age, RNA extracted and run individually on a 135k probe microarray (see (JOHNSSON *et al.* 2016)). For further details on feed and housing see (JOHNSSON *et al.* 2012b).

### Ethics Statement

The study was approved by the local Ethical Committee of the Swedish National Board for Laboratory Animals.

### Phenotyping

#### Social reinstatement assay

Also referred to as a ‘runway test’ (SUAREZ AND GALLUP 1983a), this is a basic measure of social coherence/anxiety, with stressed chicks exhibiting a stronger social cohesion response (MARIN *et al.* 2001). The individual is placed at one end of a narrow arena, with several conspecifics located at the far end. The amount of time the focal individual spends associated with the conspecifics as opposed to exploring the remainder of the arena is considered a measure of sociality/anxiety. A more social (or potentially more anxious) animal will spend more time associating with conspecifics, will approach the conspecifics more rapidly (decreased latency to approach), and will spend less time in the start zone of the arena (MARIN *et al.* 2001). Trials were replicated twice per individual, with each trial being five minutes in length. Trials were performed at three weeks of age.

Trials were performed in a 100cm × 40cm arena. The stimulus zone measured 20cm × 40cm, and was adjacent to a wire mesh compartment containing three unfamiliar conspecific birds of the same age, whilst the start zone (where the birds were placed prior to the start of the experiment) also measured 20cm × 40cm. Birds were placed in the start zone of the arena in the dark, prior to the lights being turned on and the trial beginning. Eight separate arenas were available, allowing up to eight individuals to be analysed simultaneously. Measurements were taken using the Ethovision software and continuous video recording (Noldus Information Technology, www.noldus.com). For each trial, total distance moved, velocity, proportion of time spent in the stimulus zone (adjacent to the three conspecific birds), latency to first enter the stimulus zone, length of time in the start zone (the starting position of each bird, farthest away from the stimulus zone) and the degree of meander (how directly the birds moved) was measured. Each trial was performed twice, with approximately one week between an individual’s first and second test. Individuals were immediately removed from the arena upon the completion of the test to reduce potential habituation. Repeated testing is an important method of refining the phenotypes, thereby reducing environmental noise as much as possible and allows additional variables to be extracted for each individual. In this case, average time in start and stimulus zones and average stimulus zone latency were also calculated. Furthermore, minimum and maximum time spent in the start and stimulus zones and latency to enter the stimulus zone (i.e. the higher and lower values of the two trials per individual) were also calculated. The use of these latter two values gives an estimation of the maximum (and minimum) possible anxiety values of an animal, and can potentially cut down on environmental variation, when, for example, an external stimulus disturbs the animal during one of the trials.

Correlations between the two trials were found to be extremely significant (see supplementary figure 1), indicating that the tests were strongly repeatable. A high degree of correlation was found between the social reinstatement behaviour reported here, and the open field behaviour that was also recorded for these animals (and reported in (JOHNSSON *et al.* 2016)), with 123 of the 224 pairwise correlation comparisons (these pairwise correlations were both within- and between-tests) being significant (using a 1% significance threshold, see supplementary figure 1).

### Genotyping, QTL and eQTL mapping

DNA preparation was performed by Agowa GmbH (Berlin, Germany), using standard salt extraction, with 652 SNP markers being used to generate a map of length ~92675cM, with an average marker spacing of ~ 16cM. QTL analysis was performed using the R/Qtl software package (BROMAN *et al.* 2003), with standard interval mapping and epistatic analyses performed. Interval mapping was performed using additive and additive+dominance models. In the behavioural QTL analysis batch, sex and arena were always included in the model as fixed effects, whilst body weight was included as a covariate. A Principal Component Analysis (PCA) of the first ten Principal Components (PCs) of the genotypic data were fitted to account for population substructure (see Johnsson et al(JOHNSSON *et al.* 2016) for details), with all significant PCs retained in the final model. A sex-interaction effect was added, where significant, to account for a particular QTL varying between the sexes. Digenic epistatic analysis was performed according to the guidelines given in (BROMAN AND SEN 2009). A global model incorporated standard main effects, sex interactions and epistasis was built up starting with the most significant loci and working down for each trait.

eQTL mapping was performed on the cross using R/qtl, as has already been documented previously(JOHNSSON *et al.* 2016). A local, potentially cis-acting, eQTL (defined as QTL that were located close to the target gene they affected) was called if a signal was detected in the closest flanking markers to the gene in question, to a minimum of 100 cM around the gene (i.e. 50cM upstream and downstream of the gene). The trans-eQTL scan encompassed the whole genome, and used a genome-wide empirical significance threshold. In total 535 local eQTL and 99 trans eQTL were identified previously.

### Significance thresholds

Significance thresholds for the social reinstatement QTL analysis were calculated by permutation (CHURCHILL AND DOERGE 1994; DOERGE AND CHURCHILL 1996). A suggestive significance level of a genome-wide 20% threshold was used (with this being more conservative than the standard suggestive threshold (LANDER AND KRUGLYAK 1995)). The approximate significant threshold was LOD ~4.4, whilst the suggestive threshold was ~3.6. Confidence intervals (C.I.) for each QTL were calculated with a 1.8 LOD drop method (i.e. where the LOD score on either side of the peak decreases by 1.8 LOD), with such a threshold giving an accurate 95% confidence interval for an intercross type population (MANICHAIKUL *et al.* 2006). The nearest marker to this 1.8 LOD decrease was then used to give the confidence intervals in megabases. Epistatic interactions were also assessed using a permutation threshold generated using R/qtl, with a 20% suggestive and 5% significant genomewide threshold once again used. In the case of epistatic loci, the approximate average significance threshold for pairs of loci were as follows (using the guidelines given in (BROMAN AND SEN 2009)): full model ~11, full versus one ~9, interactive ~7, additive ~7, additive versus one ~4.

### Analysis of candidate genes (eQTL genes falling within QTL intervals)

Significant social reinstatement QTL were overlapped with the previously identified eQTL, with all significant eQTL genes then being considered candidate genes for the specific social reinstatement QTL they overlapped with. To further refine these candidate genes we then modelled the gene expression value on the behavioural trait for the QTL of interest (so if an eQTL overlapped a QTL for time spent in the stimulus zone, the eQTL gene expression trait would be correlated with amount of time spent in the stimulus zone). Therefore for each eQTL overlapping a behaviour QTL, we fitted a linear model with the behaviour trait as response variable, and the expression traits as predictor, including sex and batch as factors. Weight at 42 days was included for traits where weight was used as a covariate in the QTL analysis (see Supplementary Table 1). The p-values for the regression coefficient were Bonferroni corrected for the number of uncorrelated eQTL in the QTL region. eQTL that were present within a QTL C.I. and that were also significantly correlated with the QTL trait were then considered to be candidate causative genes, and were then assessed for causality where possible (see below). One issue with this approach is that the behavioural QTL were based on up to 572 individuals, whereas the eQTL/expression phenotypes were only available for 129 individuals. Therefore the NEO and conditional QTL methods for causality testing were only applied where the behavioural QTL that a gene was potentially causative to was detectable in the smaller dataset (n=129).

Causality analysis consisted of a conditional genomic QTL scan and Structural Equation Modelling. A conditional genomic scan used combined models of the form *behaviour trait = QTL + gene expression trait + covariates + error*. These were fitted to test whether the inclusion of the gene expression trait as a covariate decreased the effect of the behaviour QTL. This effect can be used as a statistical test of a causal gene (LE BIHAN-DUVAL *et al.* 2011; LEDUC *et al.* 2011), the rationale being that if there is a causal relationship, the genotype and phenotype will be independent conditional on the gene expression. This results in the inclusion of the gene expression essentially reducing the significance of the QTL genotype factor, where the genotype is causal to the gene expression variation (in essence both the QTL genotype factor and the gene expression are explaining the same variation for the behaviour in the model, so the inclusion of both should weaken the genotypic effect.

Causality analysis was also performed using Network edge orienting software (NEO) (ATEN *et al.* 2008) to test whether the expression of correlational candidates was consistent with the transcript having a causal effect on the behaviour trait. Single marker analysis was performed with NEO fitting a causal model (marker -> expression trait -> behaviour), and three other types (reactive, confounded, collider) The NEO software evaluates the fit of the model with a χ^2^ test, a higher p-value indicating a better fit of the model. The best fitting model is chosen based on the ratio of the χ ^2^p-value to the p-value of the next best model on a logarithmic scale (base 10), called local edge orienting against the next best model (leo.nb) scores. A positive leo.nb score indicates that the causal model fits better than any competing model. Aten el al. (ATEN *et al.* 2008) use a single-marker leo.nb score of 1, corresponding to a 10-fold higher p-value of the causal model, as their threshold. They also suggest users to inspect the p-value of the causal model to make sure the fit is good (as this measures the probability another model also fits the data, the model p-value should be non-significant if the causal model fits the best). For each gene, we report leo.nb score, and p-value of the causal model.

### Global Gene Expression Correlations with Behaviour

As well as the correlations between gene expression and behaviour at the specific QTL/eQTL overlaps, a further analysis was performed for each behavioural variable whereby global gene expression (i.e. each gene in turn, covering all 36k probes) was correlated with behaviour. A standard linear model with the behaviour trait as response variable, and the expression traits as predictor, including sex and batch as factors, was used. To control for the large numbers of probes tested, significance was determined using a permutation test. For any given behaviour, the behavioural variable was permuted, with this permuted phenotype then tested against all 36k probes. The top 0.1% value was then retained from this permutation. A total of 500 permutations were performed for each behavioural measure. The top 5% of these generated an experiment-wide threshold for significance of ~4 × 10^~4^.

### Gene Network Formation

A weighted coexpression network was constructed with the WGCNA (LANGFELDER AND HORVATH 2008) package, using the gene expression data. The analysis was performed on residual values from linear models with sex and batch as predictors. Briefly, pairwise correlations are calculated between all transcripts, and a power function is applied as a soft thresholding procedure. We used the WGCNA pickSoftThreshold to choose an exponent of the power function that makes the network resemble a scale-free network. The exponent used was 5. The resulting network was divided into modules with hierarchical clustering and the Dynamic Tree Cut method of WGCNA, and the model merging function. This resulted in 26 merged modules.

We used module eigengenes, i.e. the first principal component of the expression of the transcripts in a module, to represent the gene expression profile of a module. Enrichment of Gene Ontology Biological Process, Molecular Function, and Cellular Component categories in modules were tested with DAVID (HUANG *et al.* 2008; HUANG *et al.* 2009). The cut off used for enrichment was a Benjamini-Hochberg FDR corrected p-value below 0.05.

We constructed a binary correlation network of the correlational candidate genes. We used a threshold of a pairwise Pearson correlation coefficient of 0.50. All probesets that were significantly correlated with one or more social reinstatement behaviour was included, as well as the five candidates that were identified using the combined QTL/eQTL overlap and trait correlation. We used the R package igraph (v1.0.1) for network visualization.

### eQTL clustering and sweep overlap tests

We tested for eQTL clustering and overlap between QTL and selective sweeps by comparing the number of actual overlaps to the number of overlaps in simulations of random QTL intervals. Random intervals of mean eQTL, QTL, or sweep size were placed on an interval the size of the autosomal sequenced chicken genome, correcting the length by subtracting gene-poor regions with one or fewer transcripts in a 1 Mb interval. The 95^th^ quantile of cluster size on the simulated genomes was 17. In these simulations, a set of nonredundant eQTL was used, made to represent each gene only once (created by merging eQTL for all transcripts associated with an Ensembl gene model, or that align within 1 kb of the genomic coordinates of that gene model). To define the eQTL in a cluster, we used the eQTL with most overlaps, and included all other eQTL overlapping that focal eQTL.

### Pleiotropy vs Linkage Tests

We used qxpak 5 (PEREZ-ENCISO AND MISZTAL 2011) to perform pairwise pleiotropy versus linkage tests of overlapping behaviour QTL and eQTL from clusters. A model with two separate QTL was compared to one with as single pleiotropic QTL, using a likelihood ratio test, with the χ^2^ distribution with two degrees of freedom. Nominal p-values are reported, a low p-value signifying the rejection of the pleiotropic model. Since the number of possible comparisons in eQTL clusters is prohibitively large, we only tested a few cis-eQTL at the edges, and middle of the cluster, and trans-eQTL.

## DATA AVAILABILITY

Microarray data for the chicken hypothalamus tissue are available at E-MTAB-3154 in ArrayExpress. Full genotype and phenotype data are available on figshare with the following doi: 10.6084/m9.figshare.1265060.

## RESULTS

### Social Reinstatement QTL

The different measures of sociality from the social reinstatement test were mapped individually. By combining these overlapping QTL, a total of 24 social reinstatement QTL were identified (see Supplementary Table 1), spread over 16 chromosomes. The average effect size of each QTL was 4%. The average confidence interval for the behavioural QTL was ~3Mb, indicating that the AIL generated far tighter confidence intervals than a standard F_2_ intercross. The QTL generated for social reinstatement were compared with those previously identified for open field measures. There was a strong degree of overlap between the two sets of QTL, most notably on chromosomes 1,2,4,6,7, and 10 (see supplementary table 1). Pleiotropy versus linkage tests indicated that the cluster on chromosome 2 is the result of linkage (i.e. the SR and open field QTL are separate but linked), whereas for the other clusters pleiotropy and linkage were indistinguishable, so they are either pleiotropic or too closely linked to disentangle.

### eQTL clustering

A total of 535 local eQTL and 99 trans-eQTL (from a total of 537 genes) were detected previously and have been previously reported(JOHNSSON *et al.* 2016). Very few transcripts had more than one detectable eQTL. Only 13 transcripts had both a local and a trans-eQTL, and only eight had more than one trans-eQTL. eQTL were unevenly distributed across the genome, demonstrated with a clustering test by simulation. Clusters of local eQTL on chromosomes 3, 9, 12, and 14 had cluster sizes larger than the 95^th^ quantile cluster size in our simulation (see Supplementary Table 2). The chromosome 12 cluster contains 17 trans-eQTL as well as 9 local eQTL, making it an example of a eQTL hotspot with trans effects. The clusters on chromosome 9 and 14 contain two and three trans-eQTL, respectively, while the chromosome 3 cluster is only made up of local eQTL. Supplementary Table 2 shows the composition of local and trans eQTL in clusters, and the results of pleiotropy versus linkage tests, showing some of the clusters to be more likely explained by linked but separate QTL, rather than pleiotropy (most noticeably the cluster on chromosome 9).

### eQTL candidate genes within QTL intervals

In total, eQTL for 139 genes overlap social reinstatement behaviour QTL. Correlations between eQTL expression values and the QTL trait yielded a total of 5 significant correlational candidate genes, after a Bonferroni correction for the number of uncorrelated eQTL overlapping the behaviour QTL region had been applied, shown in table 1 (see also figure 1). These represent 4 different QTL regions. Of the significant candidates, *TTRAP* was very strongly significantly correlated (P<0.001) with two of the corresponding behavioural traits, and significantly correlated (P<0.05) with three more of the corresponding traits. A total of three different genes were correlated with the QTL for minimum time spent in the start zone on chromosome 2 – *ACOT9*, *SRPX*, and *PRDX4*. In addition, this QTL also had a significant epistatic interaction with a QTL on chromosome 10. Similarly, the candidate gene *ACOT9* also has a trans eQTL that falls within this QTL interval, therefore the gene *ACOT9* not only correlates well with this QTL, but it is also present as both a local and trans eQTL at both the chromosome 2 and chromosome 10 social reinstatement QTL locations. In addition to the gene *TTRAP* for the chromosome 2 QTL at 99.3-103.3Mb, the probeset based on the EST 603866246F1 was a candidate for the social reinstatement QTL region on the first part of chromosome 2 (at 63.2-67.7Mb)

**Table 1.**
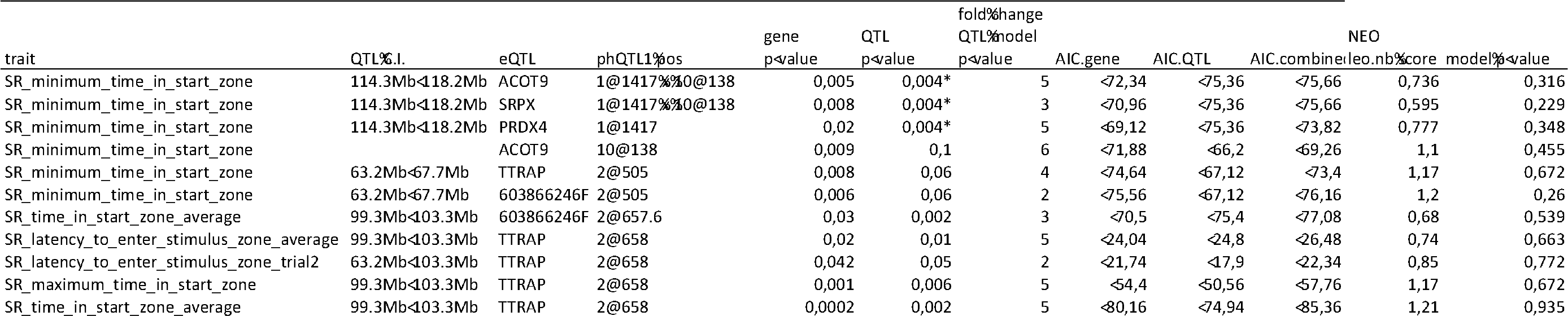
Candidate genes and causality scores.The QTL confidence interval is given in Mb, with the phenotypic (behavioural QTL) position also included in cM. Gene p-value shows the significance between gene expression and behavioural trait. QTL p-value indicates the significance between the genotype and the behavioural trait for this reduced sample size. Fold change indicates the degree to which the QTL significance drops upon inclusion of the gene expression covariate, with a higher value indicating the greater potential for causation between gene and behaviour. The AIC is also shown for the model predicting behaviour incorporating just gene expression, just the genotype and both. The NEO causality score (leo.nb) and NEO model p-value are also shown.

**Figure 1.**
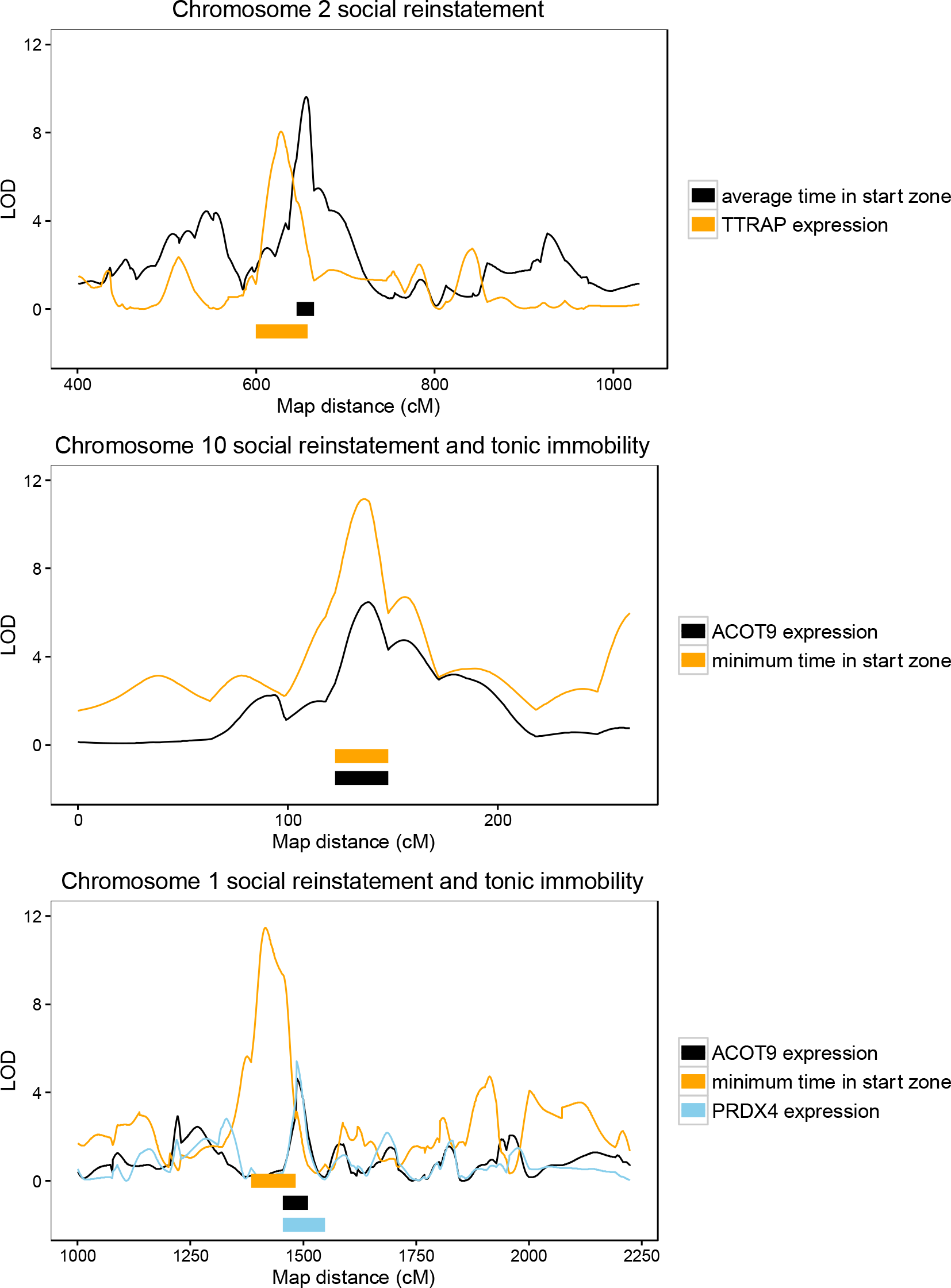
LOD profiles for *TTRAP*, *ACOT9*, *PRDX4* and their associated social reinstatement (SR) behaviours.

### Causality testing of candidate genes

Candidate genes were then dissected for causality (i.e. the likelihood they were causative QTG to their respective behavioural trait QTL) for the 5 candidate genes, see Table 1. Both NEO and conditional modelling results highlighted *ACOT9* and *PRDX4* as potentially being causal for the chromosome 1 locus for the QTL for minimum time in the start zone, whilst *SRPX* had marginally less support from the NEO analysis and is not consistent with causality at this locus. Interestingly, *ACOT9* also has a trans eQTL on chromosome 10, which also overlaps with the QTL for minimum time in the start zone on that chromosome (see table 1), with this once again receiving strong support as being causal from both NEO and the conditional modelling. The gene *TTRAP* at 99-103Mb on chromosome 2 is strongly correlated with multiple aspects of social reinstatement (see Table 1). As well as the strong correlation with these QTL (P<=0.001), there is also a large decrease in genotype significance with the gene expression covariate included in the conditional QTL model, and this gene also has the strongest support from NEO for a causal effect (leo.nb>1.2) see Table 1. In comparison, one other probe on chromosome 2 also had a correlation with time in the start zone, 603866246F1, however NEO only supported this probe for the QTL for minimum time in the start zone and not for the QTL for average time in the start zone.

### Global assessment of correlations between genes and sociality behaviour

A total of 70 genes were significantly correlated at an experiment-wide level with a variety of different measurements from the social reinstatement test (see Supplementary Table 3), with 11 of these genes correlating with multiple traits. Of these 60 genes, 11 fall within the confidence interval for one of the behavioural QTL, and are thus also putative candidate genes (*ANKRD29, CALB2, Gga.50063, HERPUD1, RDM1, SEPN1, TMEM57, TTRAP*, *TYMS, ENSGALG00000007103*, and a non-annotated probe 603602419F1), although with the exception of *TTRAP*, the behavioural trait QTL overlap was different to the actual trait that was significantly correlated with the gene (see Table 2). Interestingly, *TTRAP* is significantly correlated with both a number of different traits, and the strength of the correlation is significant even controlling for testing all genes within the genome. *TTRAP* also possessed an eQTL, whereas none of the other 11 genes did. A gene network was constructed using all 70 of the genome-wide significantly correlated genes, plus the four candidate genes generated previously (see figure 2). The gene *TTRAP* had a total of five connections, with almost all the other candidates also having high numbers of connections (see figure 2, *ANKRD29* 13 connections, *SEPN1* 18 connections, *RDM1* 0 connections, *TMEM57* 21 connections, *TYMS* 4 connections, *CALB2* 13 connections, *HERPUD1* 9 connections, *Gga.50063* 9 connections).

**Table 2.**
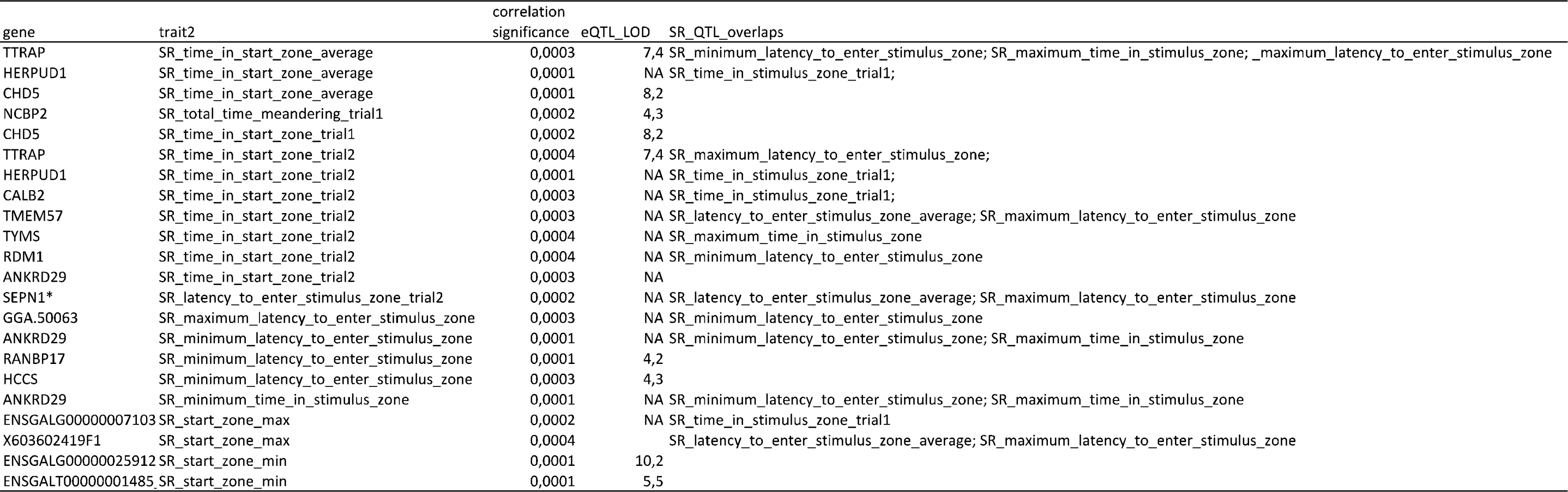
Global Gene correlation Candidates. Genes significantly correlated with behaviour at a genome-wide threshold that also possessed either an eQTL or overlapped with a behavioural QTL.

**Figure 2.**
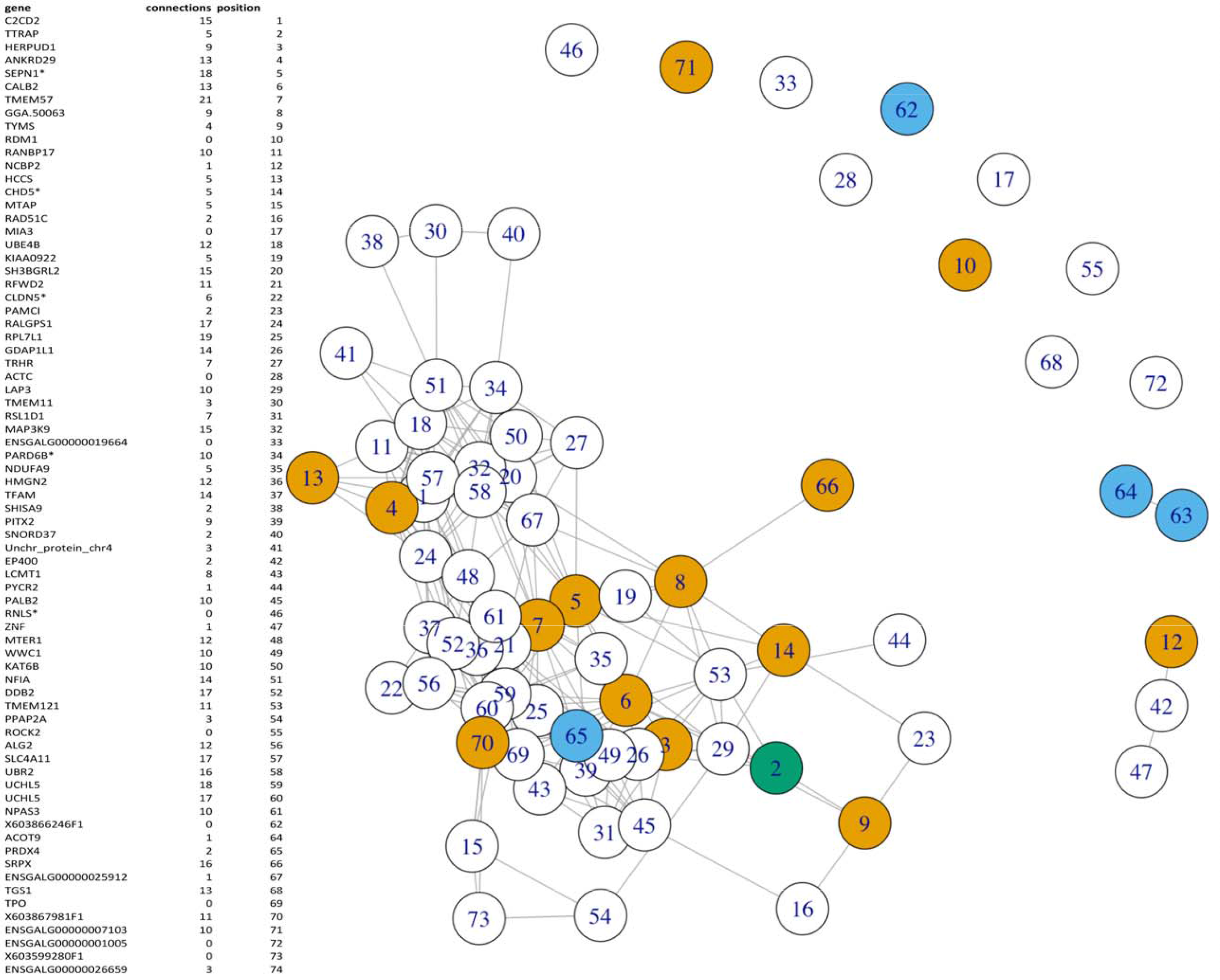
Gene network for all genes significantly correlated with SR behaviour. The number of connections each gene shares with other genes in the network is provided in the legend, as well as the key to identify specific genes within the network. Genes that were significantly correlated at the genome wide level with SR and also overlapped an SR QTL are in yellow, genes that were identified using the full overlap between an overlap of eQTL and QTL and were significantly correlated are in blue, whilst *TTRAP*, which fulfilled both these requirements is marked in green.

### Selective sweeps

Reduced heterozygosity around a locus can be a molecular signature of selection, caused by neighbouring linked variants hitch-hiking on the selected variant (MAYNARD SMITH AND HAIGH 1974) Scans for selective sweeps have been utilised to find genes selected during chicken domestication (RUBIN *et al.* 2010), though with such scans the potential phenotype that has been selected is unknown. eQTL were searched for overlaps with putative selective sweeps within domestic chicken breeds detected by Rubin et al.(RUBIN *et al.* 2010). 54 eQTL overlapped selective sweeps in layer breeds, with this being far from significant using a permutation simulation (the 5% cut-off in simulations being 92 overlapping sweeps). While individual sweeps close to eQTL may still be a signal that a particular variant affecting transcription has been selected during domestication, our data do not support a general impact of domestication-related selective sweeps on the genetic architecture of gene expression in the chicken hypothalamus.

### Module Analysis And Enrichment

The global expression values were separated into 26 gene expression modules using weighted gene co-expression analysis (LANGFELDER AND HORVATH 2008). If a gene expression module is relevant to a phenotypic trait, genes that are more central to that module should have a higher correlation to that trait in question (phrased by (GHAZALPOUR *et al.* 2006) in terms of intramodular connectivity and gene significance). The 5 significant putative QTG identified are present in a total of 4 different modules. In total, two of these genes are both well-correlated with one or more behavioural traits and have a high connectivity (correlation) with the central eigenvalue of the module (*SRPX* has a correlation coefficient of 0.68, whilst *ACOT9* has a correlation coefficient of 0.52). Two of these genes are contained in one module (orange), with the other three genes in the black, grey and yellow modules, respectively (see Supplementary figure. 1). These modules are enriched for protein phosphorylation in the cases of the black and yellow modules.

## DISCUSSION

Using a combination of QTL and eQTL mapping, followed by further correlations and causality analyses, we have identified a number of high confidence candidate genes that affect one aspect of sociality behaviour, as measured using a social reinstatement test. Of the candidates, the five genes with the highest support came from those that overlapped QTL with eQTL, showed a correlation between gene expression and behaviour and had evidence of causality (*TTRAP, PRDX4, ACOT9, SRPX* and 603866246F1). Three of these genes (*TTRAP, PRDX4* and *ACOT9*) have previously been identified as affecting either neuronal development or behaviour. Eight other candidate genes were also identified; in this case these genes were significantly correlated at a genome-wide level with an aspect of social reinstatement behaviour, and also overlapped a QTL interval. As the QTL trait was different to the correlative trait for these eight genes, causality testing could not be used, and these genes therefore have less support as candidates. We also find that eQTL are clustered in the genome, most specifically at four separate locations, where we have eQTL hotspots. These appear to contain both pleiotropic and linked loci, giving support for the idea that specific modules affect domestication (WRIGHT *et al.* 2010).

Of the five principle candidate genes that appear to affect social reinstatement, three already have some links with anxiety and stress behaviour, cognitive processes and neurogenesis and degeneration (*ACOT9*, *PRDX4*, *TTRAP)*. The best candidate, based on a wide number of measures, is the gene *TTRAP*. This was identified in both the QTL/eQTL overlap, and had the highest correlation between trait and gene expression of any of the five main candidates. In addition, the causality tests (both conditional mapping and NEO), both indicate it as being the best candidate gene. It is correlated with numerous aspects of social reinstatement behaviour, and the correlation between the trait ‘average time in the start zone’ and TTRAP expression was significant even at a genome-wide level. It also has numerous connections in the network comprising of all the candidate genes, indicating its role as a hub gene. *TTRAP* itself is a DNA phosphodiesterase, involved in DNA repair (ZENG *et al.* 2011). It has associations with early-onset Parkinson’s disease (ZUCCHELLI *et al.* 2008), and might serve a protective role against neurodegradation. Its human orthologue (*TDP2*) is required for normal neural function (Gomez-Herrero 2014). *PRDX4*, a cytoplasmic antioxidant enzyme, is linked with stress and social isolation in piglets (POLETTO *et al.* 2006), anxiety related measures in mice after treatment with paroxeline (an antidepressant) (SILLABER *et al.* 2008) and with atypical frontotemporal lobar degradation in humans (MARTINS-DE-SOUZA *et al.* 2012). Both *TTRAP* and *PRDX4* are involved in the cellular response to reactive oxygen species, and in the regulation of NFκB signalling (JIN *et al.* 1997; PYPE *et al.* 2000). NFκB signalling, in turn, is connected both to neurogenesis in response to stress (MADRIGAL *et al.* 2001; KOO *et al.* 2010), and to contextual fear memories (LUBIN AND SWEATT 2007). *ACOT9* is likely an enzyme involved in hydrolyzing acyl-coenzyme A thioesters. Loss of function of Acyl-CoA thioesterases may be involved in neuronal degradation Of the additional nine genes identified using global correlations, *CALB2* has previously been linked with schizophrenia and neuronal growth, whilst *HERPUD1* has also been linked with neuronal apoptosis and *TMEM57* has been associated with neurocognitive impairment in Alzheimer’s (LEVINE *et al.* 2013).

The genetic architecture of behaviour and its association with behavioural syndromes (where multiple character aspects are correlated across different situations) is yet to be fully explored. In this study animals have been measured for both SR and open field activity. Although the open field arena has long been considered as a measure of anxiety, it has also been posited that the test contains separate elements of fear/anxiety (leading to the inhibition of activity) and also the search for social companions (Faure 1983, Gallup and Suarez 1980, Mills 1993). This therefore implies there is a degree of overlap between the two tests. In the experiment presented here, we find that there is a strong overlap between SR and open field phenotypes, with a large number of significant correlations between the two. Our study therefore shows a stable (i.e. not disrupted by recombination) behavioural syndrome for sociality/anxiety exists in this experimental cross. Current work on behavioural syndromes uses statistical genetic correlations (DINGEMANSE *et al.* 2012) to link syndromes with their underlying genetics. Here, we show that many of the same loci affect both juvenile behavioural traits (appear pleiotropic) and exhibit a shared genetic architecture. While the species of animal and the test battery differ, our conclusions on the genetic architecture of sociality and anxiety behaviours are similar to results from mice (TURRI *et al.* 2001; HENDERSON *et al.* 2004), with independent but overlapping architectures for different behaviour tests. Like the present work, there were some shared loci, where pleiotropy cannot be excluded, and some independent loci for specific test situations.

The genetic architecture of gene expression has often been found to consist of numerous distinct ‘modules’, with these largely thought to represent pleiotropic regions. For instance, these expression modules are seen in *Drosophila* (MCGRAW *et al.* 2011), yeast (LITVIN *et al.* 2009), mice (WU *et al.* 2008) and *Arabidopsis* (WEST *et al.* 2007), amongst others. It has been suggested that this could be due to core pleiotropic effects (for example, when a single gene is perturbed, a whole raft of changes can ensue (LITVIN *et al.* 2009)). A problem is that many previous datasets use an F2 or similar study design for the eQTL analysis, or if RILs are used, the sample sizes (in terms of numbers of different lines) are very low. In the study shown here, the narrow intervals generated by the AIL make it possible to start to disentangle linked effects from pleiotropy. Four distinct modules of eQTL were seen in this analysis, on chromosomes 3, 9, 12 and 14. Pleiotropy versus linkage tests reveal that not all these eQTL modules are pleiotropic. For example, the eQTL module on chromosome 9 (consisting of 26 eQTL) contains a pleiotropic ‘core’ of eQTL (or at least a core where pleiotropy is indistinguishable from linkage with this being surrounded by linked, separate, loci. Similarly, the pleiotropic hotspot on chromosome 14 has a pleiotropic core of local eQTL, but the two trans-eQTL that are also part of this module are linked. Finally, one of the loci on the chromosome 3 hotspot shows some evidence of linkage as well. Conversely, a more standard putatively fully pleiotropic hotspot is also seen on chromosome 12. These combinations of pleiotropic cores, surrounded by satellite linked loci mirror what has been seen for domestication, using behavioural, morphological and life-history traits (WRIGHT *et al.* 2010). It therefore appears that whilst pleiotropy is indeed present for such domestication traits, these clustered regions affecting domestication also contain linked effects. Similarly, the lack of power to detect weak trans effects is a problem in eQTL studies, and probably explains to a large extent the lack of trans-eQTL. However, our results suggest that the major gene regulatory changes during domestication are mostly local changes that affect neighbouring genes locally or a few genes in trans.

The potential for domestication to lead to the presence of selective sweeps around fixed loci could lead to rapid identification of the causative mutation affecting such behavioural traits. In strong contrast to a sexual ornament (comb mass), which showed a strongly significant enrichment for selective sweeps present in comb mass QTL (JOHNSSON *et al.* 2012a), neither behavioural nor eQTL here showed such enrichment. This potentially indicates that whereas certain alleles for morphological traits have been fixed in many different domestic populations, this is not the case for behavioural traits. This would imply that such behavioural alleles are rather population specific, potentially due to the different population history of chicken breeds causing different subsets of the many loci for fearful behaviour being fixed in different breeds. Alternatively, this could indicate that for behavioural variation, polygenic adaptation through changes in gene frequency may have a greater effect. In this instance many more loci may be present, and these would be ‘hidden’ from standard selective sweep detection as a result, with the standard sweep indicators no longer helpful for such alleles (PRITCHARD AND DI RIENZO 2010).

In conclusion, a combination of behaviour QTL mapping and transcriptome-wide eQTL mapping identify 5 primary candidates for social behaviour in the chicken. The overlap between QTL and eQTL, behaviour–gene expression correlations, and structural equations modelling support them as quantitative trait genes. Most of them are previously known to affect behaviour or nervous system function, however this is the first time for at least four of the genes that they have been implicated in sociality. The advanced intercross design gives us high resolution to detect multiple QTL of modest effect, and demonstrate that overlapping loci underlie correlated behaviours at the phenotypic level, with evidence of a modular basis for a behavioural syndrome, with both pleiotropic and linkage effects.

---

Supplementary Figure 1. Heatmap of correlations between Social Reinstatement and Open field behaviour. Values above the diagonal are the Pearson correlation coefficient, whilst values below the diagonal indicate significance (using an FDR adjust p <0.01).

Supplementary Table 1. All detected social reinstatement (SR) QTL with previously identified open field QTL. Chromosome, position (in cM), additive and dominance effect, r squared, lower and upper confidence interval (CI)in cM, nearest upstream and downstream marker to the C.I., covariates incorporated in the model and any interactions in the model are all included.

Supplementary Table 2. Pleiotropy vs linkage tests

Supplementary Table 3. All genes significantly correlated with behaviour at a genome-wide threshold.

